# Interpretable Prediction of mRNA Abundance from Promoter Sequence using Contextual Regression Models

**DOI:** 10.1101/2022.08.27.505543

**Authors:** Song Wang, Wei Wang

## Abstract

While machine learning models have been successfully applied to predicting gene expression from promoter sequences, it remains a great challenge to derive intuitive interpretation of the model and reveal DNA motif grammar such as motif cooperation and distance constraint between motif sites. Previous interpretation approaches are often time-consuming or hard to learn the combinatory rules. In this work, we designed interpretable neural network models to predict the mRNA expression levels from DNA sequences. By applying the Contextual Regression framework we developed, we extracted weighted features to cluster samples into different groups, which have different gene expression levels. We performed motif analysis in each cluster and found motifs with active or repressive regulation on gene expression as well as motif combination grammars including several motif communities and distance constraints between cooperative motifs.

## Introduction

Promoters are critical for regulating gene expressions. The promoter sequences define the strength, number and position of transcription factor (TF) binding sites, which in turn regulate the transcriptional levels of mRNA (Huminiecki and Horbańczuk, 2017; Sanchez, et al., 2011; Tang, et al., 2020). In eukaryotic cells, a promoter sequence can be divided into three regions based on the distance from the transcription start site (TSS): core promoter, proximal promoter, and distal promoter (Aysha, et al., 2018). The core promoter contains TSS, key DNA sequence elements such as TATA box and downstream promoter element (DPE) (Ngoc, et al., 2020). The proximal promoter is upstream from the core promoter, where transcription factors (TFs) mainly bind to (Huminiecki and Horbańczuk, 2017). The distal promoter is further upstream from the proximal promoter that often contains weak TF binding sites (Thonpho, et al., 2013). Uncovering the information encoded in the promoter sequences crucial for transcriptional regulation and unraveling the regulatory rules between gene expression and DNA sequences remain as an important problem.

Previous studies have analyzed the type, number, location, orientation of TF motifs, combinatorial strategies of different TF motifs, and the surrounding sequences of the TF binding sites in the promoters (Cheng, et al., 2014; de Boer, et al., 2020; de Jongh, et al., 2020; Fan, et al., 2021; Haberle, et al., 2019; King, et al., 2020; Levo, et al., 2017; Meyer, et al., 2013; Perez-Pinera, et al., 2013; Sinha, et al., 2008; van Arensbergen, et al., 2017; Weingarten-Gabbay and Segal, 2014; White, et al., 2016; Whitfield, et al., 2012; Won, et al., 2008; Xiang, et al., 2020; Xie, et al., 2013). For example, several known sequence motifs such as the TATA box (TATAWAAR) and the initiator sequence (YYANWYY in human) located at the fixed position in the core promoter region have been discovered (Juven-Gershon and Kadonaga, 2010). Distance constraints between motif combinations have also emerged from analyzing various TF binding sites. For example, the ETS:IRF composite element (EICE) prefers two nucleotide-long spacers and ETS:IRF response element (EIRE) prefers three nucleotide-long spacers (Nagy and Nagy, 2020). Furthermore, additional insights are obtained from recent efforts on generating millions of synthetic promoter sequences and measuring their impacts on gene expressions (de Boer, et al., 2020; de Jongh, et al., 2020; Ngoc, et al., 2020; Vaishnav, et al., 2022). Despite these progresses, there remain great challenges of uncovering the regulatory grammar encoded in the promoter sequences.

Several studies have been reported to predict the transcription strength or mRNA level from promoter or core promoter sequences by using various deep learning models (Agarwal and Shendure, 2020; Ngoc, et al., 2020; Vaishnav, et al., 2022; Zheng, et al., 2021; Zrimec, et al., 2020). They found that the gene expression levels are controlled by the entire gene regulatory structure and specific combination of regulatory elements rather than single motifs or genomic regions (Zrimec, et al., 2020). For example, different combinations of promoter motifs lead to different expression levels (Andersson and Sandelin, 2020; Aysha, et al., 2018; de Boer, et al., 2020). However, the models for studying the regulatory rules are usually complex. Due to the black-box nature of deep learning (de Jongh, et al., 2020), these models still lack a clear interpretation of the promoter architecture, such as the motif location and identity in the promoter regions that have the highest active or repressive effects on gene expression. Furthermore, they cannot reveal motif grammar such as the interactions of the TF binding sites, motif community, and coregulation effects of the motifs. A few interpretation approaches were reported, such as the perturbation impact map and saliency map (Talukder, et al., 2021), but they are often time-consuming or hard to learn the combinatory rules (de Jongh, et al., 2020).

We have developed a framework called contextual regression to interpret nonlinear models (Liu, et al., 2019; Liu and Wang, 2017). Here, we designed a workflow implementing this framework to predict the mRNA levels from the promoter sequences. The model can uncover the most predictive features by using the dot product of the extracted features of DNA segments with a certain length and the context weight. The weighted features allow to cluster the promoter sequences into groups with different mRNA levels. By analyzing the extracted features in each group, we uncovered the DNA segments with the active or repressive effects on expression, and found the enriched motifs in these segments. This workflow is flexible to be applied to seven different promoter sequence ranges (10.5kb, 800bp, 400bp, 200bp, 100bp, 50bp, and 19bp around TSS) with increasing resolution. Starting from the analysis of 10.5kb promoter sequences around TSS, we found that there are several discrete sequence stripes showing higher contribution to the gene expression level. By comparing the co-occurrence locations of discovered motifs, we also found the motif grammars including the motif communities and motif pairs with specific distance constraints. Then, we analyzed regions of 800bp, 400bp, 200bp, 100bp and 50bp around TSS to increase resolution of locating the most predictive segments and motifs. Lastly, we focused on the downstream promoter region (DPR, +17∼+35bp around TSS) sequences and elucidated the differences between synthetic and genomic sequences on regulating gene expression as well as identified several bases strongly preferring guanine in highly expressed genes.

## Results

### Predicting gene expression using promoter sequences

Using the contextual regression model (Liu, et al., 2019; Liu and Wang, 2017), we firstly extracted features from the 10kbp promoter sequences. We found that the promoter sequences are not equally important for predicting gene expressions and particular locations in the upstream of TSS are crucial for regulating transcription. Distinction between the highly-expressed and the lowly-expressed genes largely come from sequences close to TSS. To study the sequence features at finer resolutions, we trained additional models on the sequences of 800bp ∼ 50bp around TSS and also the downstream promoter region (DPR, +17∼+35bp around TSS).

We first trained a contextual regression model (Liu, et al., 2019; Liu and Wang, 2017), referred to as CR-1, to predict gene expression levels using promoter sequences that span from -7kb to +3.5kb around the TSS (Fig. 1A). The model was composed of three convolutional blocks, followed by two fully connected blocks and one fully connected layer. The convolutional filter lengths were 7, 7, 7, and the convolutional filter numbers were 32, 16, 8. The strides of max pooling layers were 50, 2, 2, which resulted in 200bp bins in the extracted feature vector. The numbers of neurons in the fully connected layers were 64, 2, 410 (52 bins × 8 filters). To avoid overfitting, a dropout layer was added after the first two fully connected layers and the dropout probability was set to 0.00099. In the third fully connected layer, we applied L1 regularization on the weight with a penalty coefficient 0.0001 to make the weighted features more interpretable. The model performed well as the predicted and measured values are close to each other along the diagonal line in the scatter plots (Fig. 1B and 1C). The Pearson correlations for the training and testing datasets were close to each other (0.736 and 0.720, respectively), which suggests no overfitting. These correlation values are also comparable with the Xpresso results (Agarwal and Shendure, 2020) (Supplemental Table S3).

**Fig. 1.**
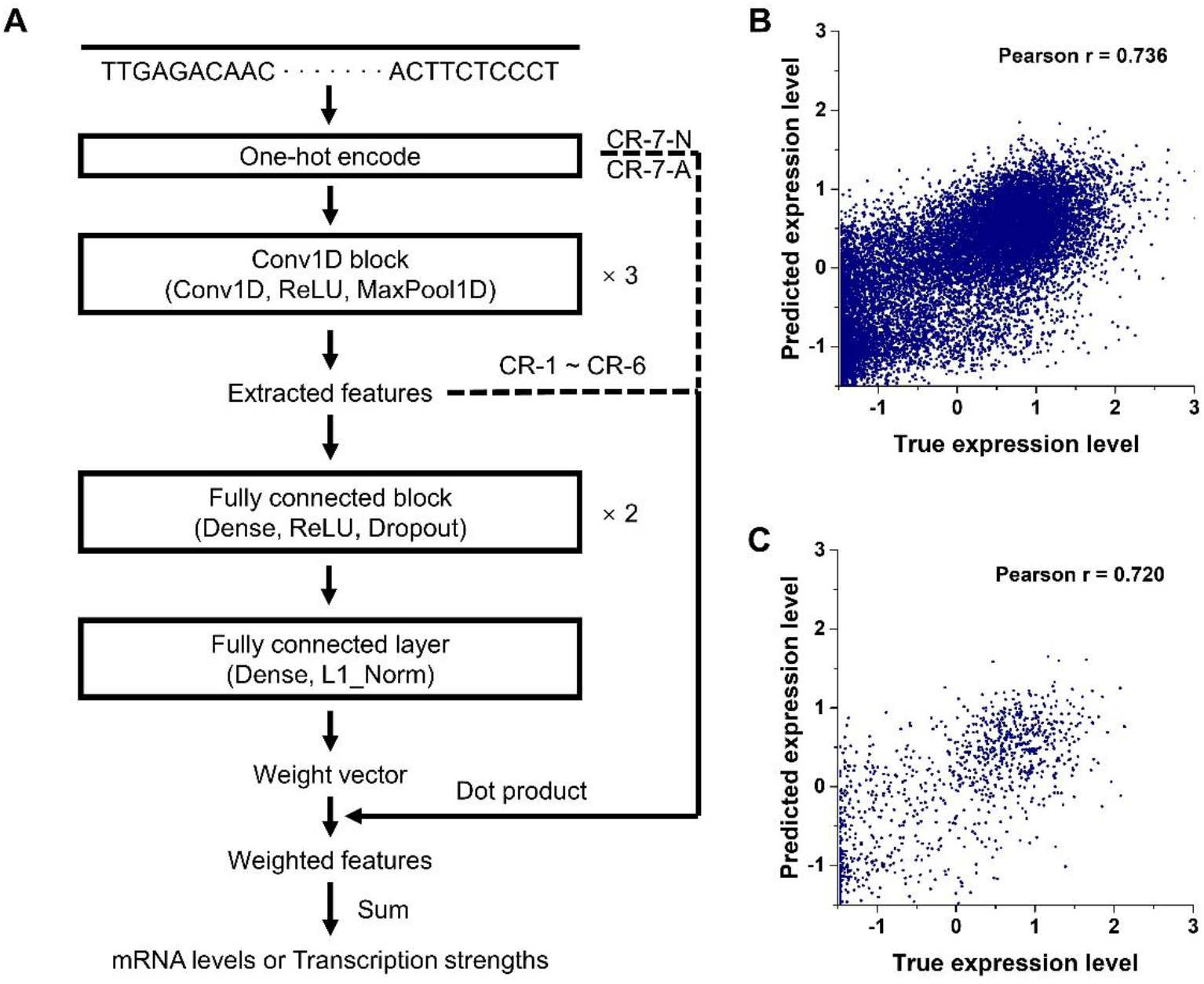
(A) The contextual regression model structure. (B) Prediction performance of CR-1 for the training dataset. (C) Prediction performance of CR-1 for the testing dataset.

The model CR-1 successfully uncovered the regions with positive and negative contextual weights (indicating active or regressive effects on gene expression) that are most predictive of gene expression (see below and Fig. 2). Interestingly while not surprisingly, the contextual weights of the highly and lowly expressed genes are most distinct on the region next to TSS. As the CR-1 model was 200-bp resolution considering sequence information extraction efficiency and prediction ability, we zoomed into the -400bp ∼ +400bp sequences corresponding to the 33rd to 36th bins to retrain the contextual regression model with higher resolution (referred to as CR-2 model). We changed a few hyperparameters as listed below from CR-1 to CR-2. The strides of max pooling layers were 4, 2, 2, which resulted in 16bp-bins in the extracted feature vector. The numbers of neurons in the fully connected layers were 8, 2, 400 (50 bins × 8 filters). We also checked the prediction performance of model CR-2. As shown in Supplementary Fig. S1, the Pearson correlations for the training and testing datasets are 0.709 and 0.688 respectively. Since the sequences used in CR-2 are much shorter than CR-1, it is reasonable that the correlation values in CR-2 are slightly lower than CR-1. Such a minor difference suggests that the sequences around the TSS heavily govern the gene expression.

**Fig. 2.**
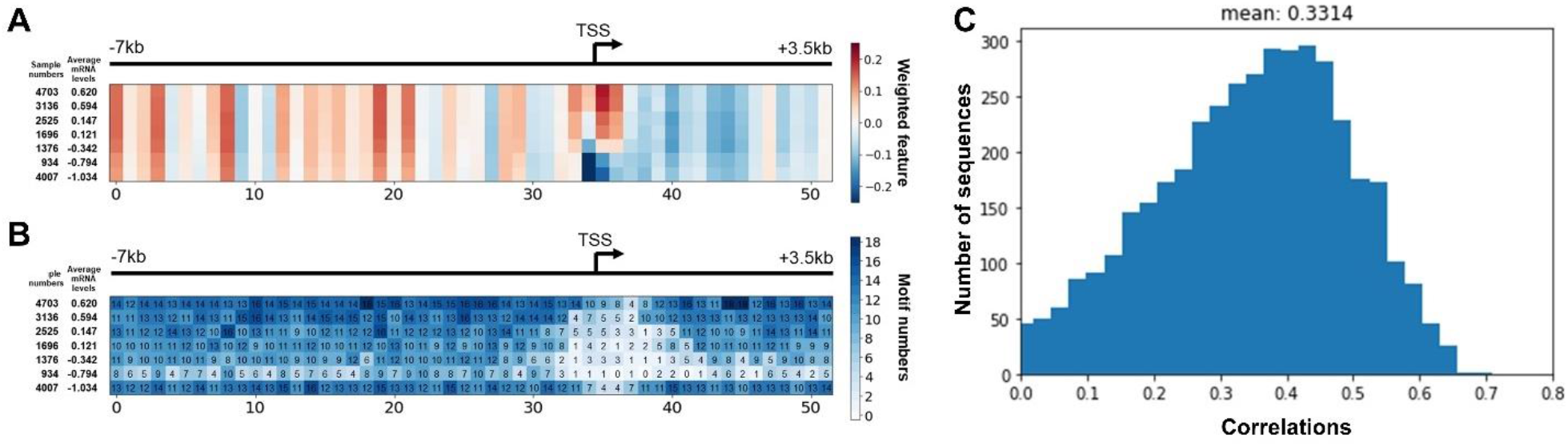
(A) The average weighted features of 7 groups of samples. Red and blue colors represent positive and negative contributions to the prediction, respectively. The TSS is located in the 35^th^ bin (bin index from 0 to 51). (B) The motif numbers in each group and each bin position. In each figure, the top shows the sequence range and TSS location. Left shows the sample numbers and the average mRNA levels (y^ = log_10_(y + 0.1)). The groups are ordered by the descending order of average mRNA levels. (C) The distribution of correlations between the weighted features and the non-linear combination of CG content and H3K27ac signal.

To further improve the resolution of defining the sequence contribution to gene expression with a focus on the region around TSS, we trained another four models for the sequences with lengths of 400bp, 200bp, 100bp, and 50bp around TSS. The last model was able to locate every base in its 50-bin length weighted feature layer. As shown in Supplementary Fig. S2-S5, the Pearson correlation of CR-3 (−200bp ∼ +200bp around TSS) was 0.696, only slightly decreasing from CR-1 and comparable with CR-2, while CR-4 to CR-6 with further decreased prediction performance with shorter sequence length. This observation suggested -200bp ∼ +200bp sequences around TSS likely contain the majority of the regulatory information of gene expression.

We also analyzed the proximal downstream region around TSS (i.e. the DPR regions which is located in the 26th bin in the model CR-2) and focused on the sequence of +17bp to +35bp relevant to TSS. This region has been shown to be crucial for transcription by, for example, analyzing gene expressions controlled by random sequences (Ngoc, et al., 2020). We trained another contextual regression model CR-7-A for this region on the synthetic dataset (Ngoc, et al., 2020). The model structure was slightly adjusted (Supplementary Fig. S6). The weight layer was multiplied with the one-hot-encoded layer instead of the output after three convolutional layers, which let each node in the weighted feature matrix correspond to each base. The strides of max pooling layers were 2, 2, 2. The numbers of neurons in the fully connected layers were 16, 2, 76. The Pearson correlations for the training and testing datasets are 0.896 and 0.836 respectively, indicating that the model successfully captured the regulatory relationship between sequences and gene expressions (Supplementary Fig. S7). As a comparation, we also used genomic sequences to train another model CR-7-N for the same region. The Pearson correlations for the training and testing datasets are only 0.486 and 0.483 respectively (Supplementary Fig. S8). This observation suggests that the genomic sequences of +17bp to +35bp only account for a small portion of random sequences and are insufficient for regulating transcription, highlighting the importance of sequences outside this region in the natural promoters in precise control of gene expressions.

We further analyzed the base preferences of the genomic (CR-7-N) and synthetic (CR-7-A) datasets in DPR. As shown in Supplementary Fig. S15, we counted the base frequencies of sequences in DPR and compared them in the two groups with the highest (red line) and lowest (blue line) one percent of expression level. Overall, the genomic sequences have more G and C than A and T consistently in all positions while the synthetic sequences show larger fluctuation and some positions such as +30 have more A/T than C/G. Another striking observation is that G has a much higher percentage in the synthetic than in the genomic sequences. Furthermore, the percentage profile of each base is also quite different between the genomic and synthetic sequences. These observations suggest that the genomic DPR sequences lack the additional features included in the synthetic sequences to regulate gene expression, which is consistent with that CR-7-N could not accurately predict gene expression levels using the genomic DPR sequences.

### Promoter sequences are not equally important for regulating gene expression

The weighted feature layer of CR-1 includes 410 neurons, corresponding to 52 bins, i.e. 8 neurons per bin. Before visualizing the features, the 8 weighted features belonging to each bin were summed together, resulting in 52-bin long weighted feature vector for each 10,500-bp promoter sequence. The similar feature processing procedures were performed for CR-2 ∼ CR-6, leading to 50-bin long weighted feature vectors for each 800bp ∼ 50bp promoter sequence. The above six vectors were concatenated together resulting in 302-bin length weighted feature vector for each sequence. Next, we clustered all promoters into 7 groups based on their 302-bin weighted feature vectors using the Ward linkage criterion (Supplementary Fig. S9). Then, we calculated the average expression levels (Supplementary Fig. S10) and averaged weight features for the samples in each group (Fig. 2A, Supplementary Fig. S11A, and Supplementary Fig. S12).

Clearly, the promoter sequences are not equally predictive of gene expression (Fig. 2A) and several upstream regions show strong activating effects on transcription (positive contextual weights), such as 0^th^, 3^rd^, 8^th^, 19^th^, and 2^st^ bins. These stripes of high contextual weights suggest that these locations have more contribution to the gene expression regulation than the other locations. The most prominent distinction between the highly and lowly expressed genes is in the bins next to TSS, consistent with the literature that these core promoter regions are crucial for transcriptional regulation. For the model with finer resolutions (Supplementary Fig. S11A and Supplementary Fig. S12), different promoter regions also exhibit different contributions to gene expression, which helps us gradually zoom in to key regions and eventually reach base-pair resolution. In the DPR sequences, most of positions show positive contributions especially for +20, reflecting its important ability of gene expression regulation.

To find what sequence and chromatin features may contribute to the striping patterns of regulatory importance in promoters, we calculated the correlations between each sequence’s weighted features and several simple sequence features including CG content, H3K27ac, H3K4me1, and H3K4me3. We used the random forest to perform non-linear regressions and found the combination of CG content and H3K27ac from H1-hESC cell line had the highest correlation to the weighted feature (Fig. 2C).

### Identification of motifs and regulatory grammar of the promoters

The largest contributions came from the bins around the TSS, ranging from activating to repressing transcription consistent with the gene expression levels. This result indicates that the major sequence difference between variant groups is largely around the TSS including the core promoter region. We used the motif finding software STREME (Bailey, 2021) to find motifs in each group and each 200bp bin defined in model CR-1 (200bp provided a sufficient segment for motif finding) with a p-value cutoff of 0.001 (Fig. 2B). We found 4003 motifs in total. To remove redundant motifs, we used Tomtom (Gupta, et al., 2007) to determine the similarities between all pairs of motifs (Fig. 3A and Fig. 3B). Using a p-value cutoff of 10^−5^, we extracted 87 unique motifs.

**Fig. 3.**
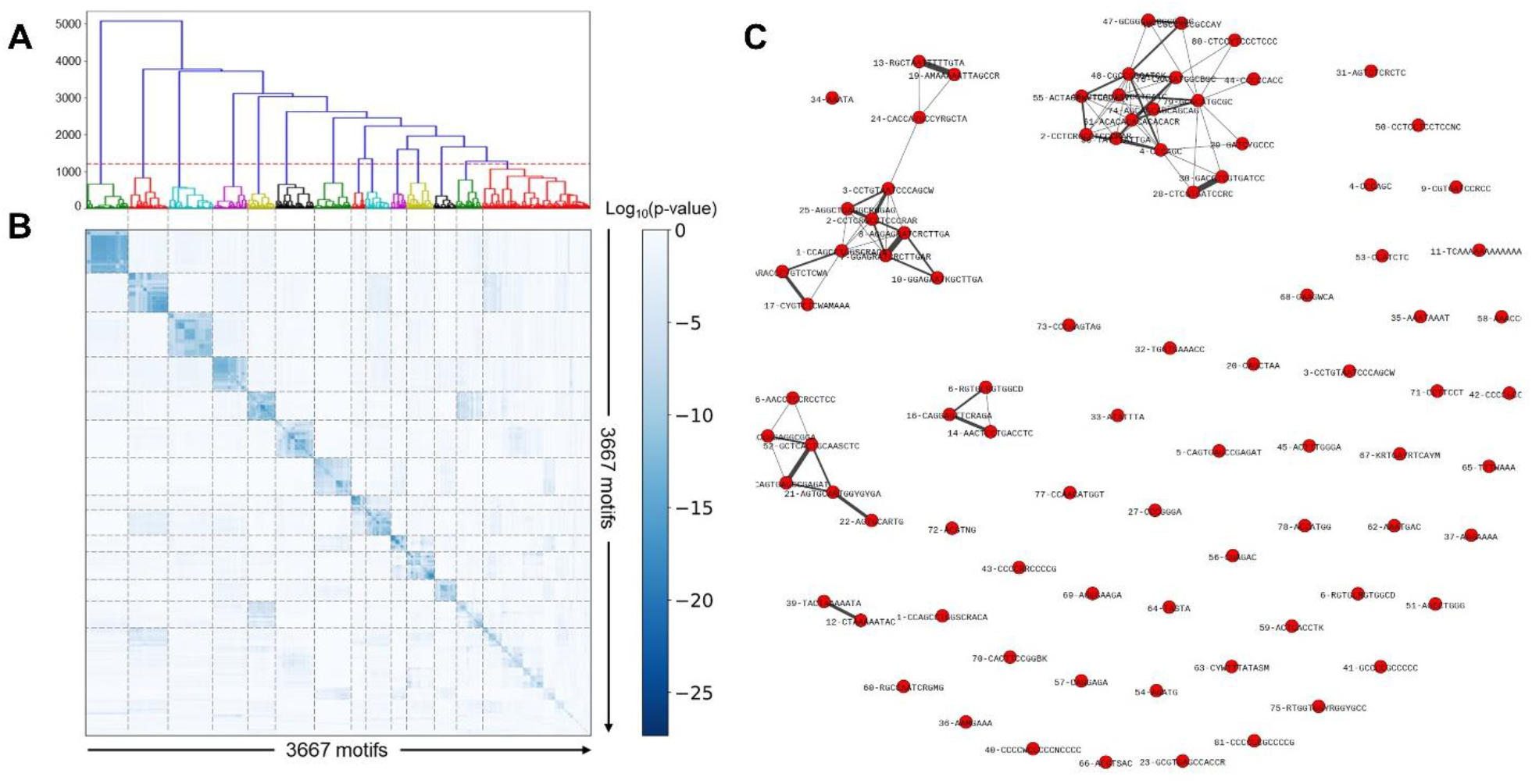
(A) The hierarchical clustering of similarity vectors of 3667 motifs by using the Ward variance minimization algorithm. The red dashed line is the distance threshold of 1200. (B) The similarities between all pairs of motifs were calculated using Tomtom. (C) The motif communities for the samples in the group with the highest expression level. The lay-out was generated using the Fruchterman-Reingold force-directed algorithm and the width of the edge represents the cosine similarity score.

We compared these 87 motifs with the known ones in HOCOMOCO v11 (Kulakovskiy, et al., 2018) as well as the DNA motifs that are associated with histone modifications (361 motifs) (Ngo, et al., 2019), and DNA methylation (313 motifs) (Wang, et al., 2019) using Tomtom with an E-value cutoff of 0.05 (Supplemental Table S4). The 28 matched known motifs include TFs, such as well-known promoter binding factors of TBP and TAF1 (Anandapadamanaban, et al., 2013) as well as SP4 and EGR1, which have been reported to regulate gene expression by binding to the CG rich promoters (Maag, et al., 2017). Surprisingly, the majority (64 motifs) of the 87 motifs matched with motifs associated with histone modifications (50 motifs) (Ngo, et al., 2019) and DNA methylation (39 motifs) (Wang, et al., 2019) (see details in Supplemental Table S4). The matched histone motifs are associated with histone modifications of H3K27ac, H3K27me3, H3K36me3, H3K4me1, H3K4me3, and H3K9me3. This observation highlights the importance of epigenetic modifications and the factors involved in establishing or maintaining these modifications on regulating gene expression.

Next, we investigated whether there is any distance constraint between the co-occurring motifs. We used FIMO (Grant, et al., 2011) to find all motifs’ occurrence sites and calculated the cosine similarities with different lags for all pairs of motifs (Supplementary Fig. S13). For a pair of motifs, if their average cosine similarity is larger than 0.7 and co-occurrence frequency is larger than a quarter of the total sequence numbers in the group, they were considered to tend to co-occur. Fig. 3C shows that these motifs form three large communities in which the motifs are densely connected to one another.

For the 82 pairs of motifs that show high cosine similarity (78 are within the same community shown in Fig. 3C), we checked whether they tend to co-occur with certain distance constraint. Fig. 4A shows that more than half of them prefer to co-occur in the same 200-bp bins and interestingly these bins avoid the region around TSS (Supplementary Fig. S14). To further reveal distance constraint rules on base-pair resolution, we checked 45 motif pairs preferring to co-occur in the same 200bp bins. We found that most of the motif pairs do show preferred distance spacing between their occurrences (Fig. 4B). For example, the distance spacer between motif 21 (AGTGCARTGGYGYGA) and motif 52 (GCTCACTGCAASCTC) prefers to be -20bp (accounting for 96% of all the occurrences). Another example is motif 1 (CCAGCCTGGSCRACA) and motif 2 (CCTCRGCCTCCCRAR) that mostly prefer to be 47bp or 45bp apart (accounting for 42% of all the pairs).

**Fig. 4.**
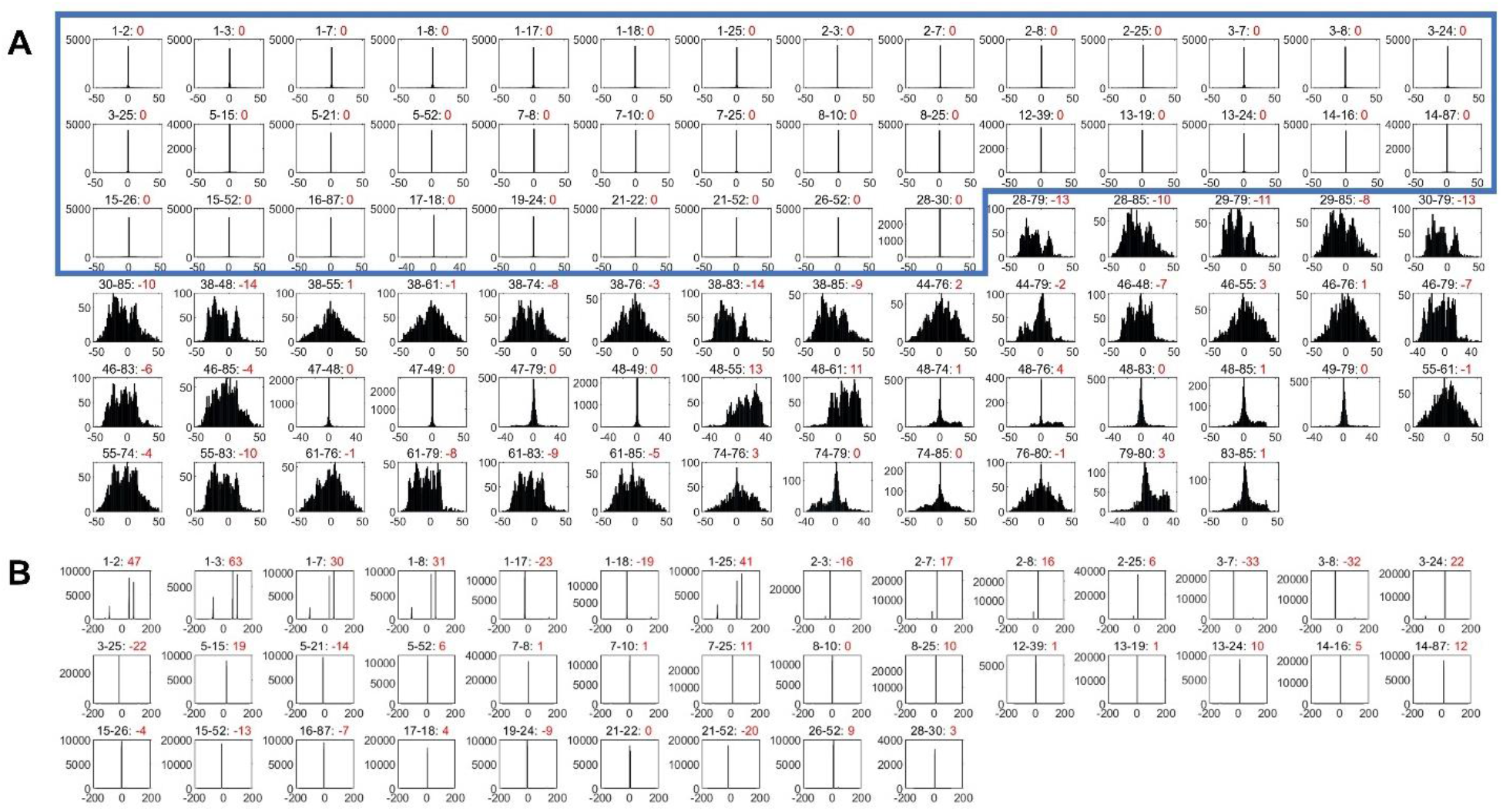
(A) The lag distribution for the 82 co-occurring motif pairs. The numbers in red are the median values of the lag. The blue box includes the 37 motif pairs that prefer to co-occur in the same 200bp-bin. (B) The distance constraint for the 37 co-occurring motif pairs. The numbers in red are the median values of distances (bp).

## Discussion

In this study, we trained interpretable neural network models based on the contextual regression framework, which can not only predict the gene expression levels from the DNA sequences but also reveal the key features by using the contextual weight. The first model CR-1 with resolution of 200bp found several stripes which provide the major contributions to the gene expression levels with the active or repressive effects. These stripes are related to the non-linear combination of CpG signals and H3K27ac signals. In particular, the regions around TSS show the most distinction between highly and lowly expressed genes. We thus built higher resolution models (CR-2 to CR-6) zoomed into the regions around TSS to illustrate the most important contributing sequences. A repeating observation is that the promoter sequences are not equally important for gene expression, suggesting the importance of the underlying promoter sequences in regulating transcription.

Furthermore, by combining CR-1 to CR-6 models with increasing resolution, we found the promoter regions around TSS have the most distinction between highly and lowly expression genes. Using the contextual weight profiles, we could cluster all the genes into 7 groups with expression levels ranging from high to low, suggesting that the contextual weight reflect the sequence features associated with transcriptional regulation.

An interesting observation is that the CR-7-N model trained on the genomic DPR regions (+17bp to +35bp relative to TSS) could not predict gene expression well while the gene expression levels of the synthetic DPR sequences could be accurately predicted, suggesting the necessity of promoter sequences beyond the DPR in the genomic promoters on regulating transcription. Our analysis revealed that the genomic and synthetic sequences differ in multiple ways such as CG content and preference of certain bases in some positions, suggesting possible features lacked in the genomic DPRs for controlling gene expression.

We next discovered 87 unique motifs important for predicting gene expression, among which 28 are matched with known TF motifs including those important for transcription such as TBP and TAF1. Interestingly, 64 out of the 87 motifs (74%) are matched with epigenetic motifs including 50 matched with histone associated motifs, and 39 with DNA methylation associated motifs. While the epigenetic motifs are supposed to be associated with establishing or maintaining epigenetic modifications and their importance in regulating gene expression is not unexpected, their dominance in the 87 unique motifs is still surprising and encourages future studies of the underlying mechanisms.

Our analysis also revealed the motif combination grammars including three motif communities and distance constraint rules. The three communities represent possible collaboration between a set of regulatory proteins. There are 82 pairs of motifs having high co-occurrence frequency, and about half of them have preferred distance spacing in the same 200-bp bins. This observation indicates strong cooperation between the regulatory proteins binding to the promoters.

## Methods

The promoter sequences and corresponding mRNA expression levels were downloaded from (Agarwal and Shendure, 2020). The dataset contains 18,377 genes in 56 human cell types generated by the NIH Roadmap Epigenomics Consortium. Following the same procedure as in (Agarwal and Shendure, 2020), the median mRNA expression levels across 56 cell types were used for prediction because mRNA expression levels are highly correlated (average correlation of 0.78) between different cell types (Agarwal and Shendure, 2020). For training the contextual regression models, we selected 1000 genes as the test dataset and another 1000 genes as the independent validation dataset. The remaining 16377 genes were used as a training dataset. We performed 10 times of cross validation by randomly partitioning the dataset to verify the consistency of our models (Supplemental Table S1). The synthesized DPR sequences in the core promoter region and their corresponding transcriptional strength were downloaded from (Ngoc, et al., 2020). Among the 468,069 sequences with measured transcriptional level, 7,500 sequences were selected as the test dataset, 20,000 sequences as the validation dataset, and 180,000 sequences as the training dataset.

We applied the same model structure with different fine-tuned hyperparameters to seven ranges of DNA sequences: the first one (the CR-1 model) for the 10.5kbp sequences around the TSS (−7kbp, +3.5kbp); the second one (CR-2) for the -400bp to +400bp sequences around the TSS; the third one (CR-3) for the -200bp to +200bp sequences around the TSS; the fourth one (CR-4) for the -144bp to +56bp sequences around the TSS; the fifth one (CR-5) for the -112bp to -12bp sequences around the TSS; the sixth one (CR-6) for the -92bp to -42bp sequences around the TSS; and the seventh one (CR-7-N for genomic sequences and CR-7-A for synthesized sequences) for DPR in core promoters that are +17∼+35bp relevant to the TSS. In each contextual regression model, the sequence features were extracted by a series of convolutional layers and subsequently several fully connected layers were applied to generate a weight vector with the same dimension of the sequence features. Finally, the model output was obtained by summing the dot product of the feature (CR-1 to CR-6) or one-hot-encoded vector (CR-7-N and CR-7-A) vector and the weight vector (Fig. 1A). The models were built and trained on TensorFlow 1.15.2 (Abadi, et al., 2016) and Keras 2.2.4 (Chollet, 2015). The hyperparameters were slightly adjusted from those used in (Agarwal and Shendure, 2020) for a specific dataset and model (Supplemental Table S2). The initial parameters were generated by the Glorot normal initializer (Glorot and Bengio, 2010). The SGD optimizer was used to optimize the parameters with a learning rate of 0.0005 and momentum of 0.9.

## Data access

The code and intermediate analysis data sets generated in this study are available at GitHub (https://github.com/Wang-lab-UCSD/CR-for-Promoter) and as Supplemental File.

## Acknowledgements

We would like to thank Mr. Chengyu Liu, Dr. Lina Zheng, and Dr. Richard Ainsworth for their helpful comments and discussions. This work was partially supported by NIH (R01HG009626 to W.W.).

