## Supplementary Materials for "Interpretable Prediction of mRNA Abundance from Promoter Sequence using Contextual Regression Models"

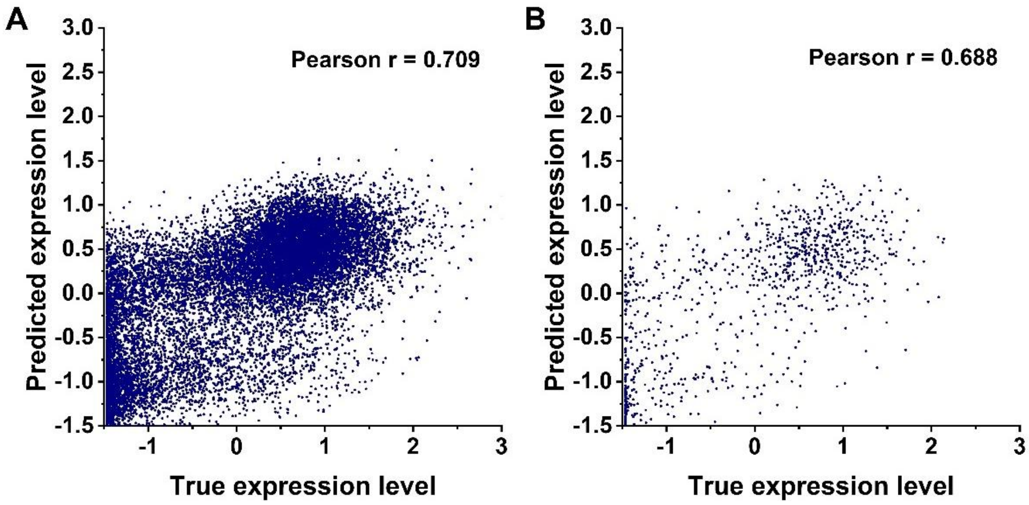


**Supplementary Fig. S1.** Prediction performance of CR-2 for (A) training dataset and (B) testing dataset.

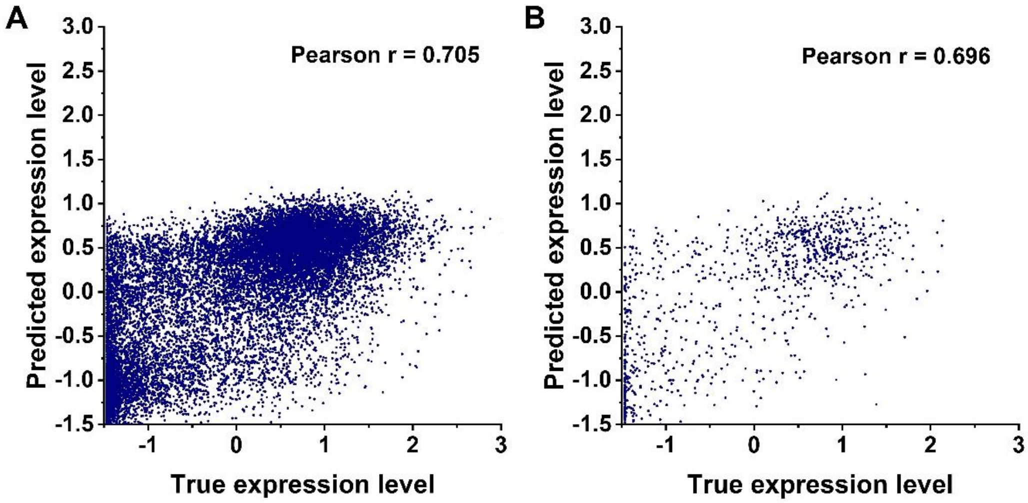


**Supplementary Fig. S2.** Prediction performance of CR-3 for (A) training dataset and (B) testing dataset.

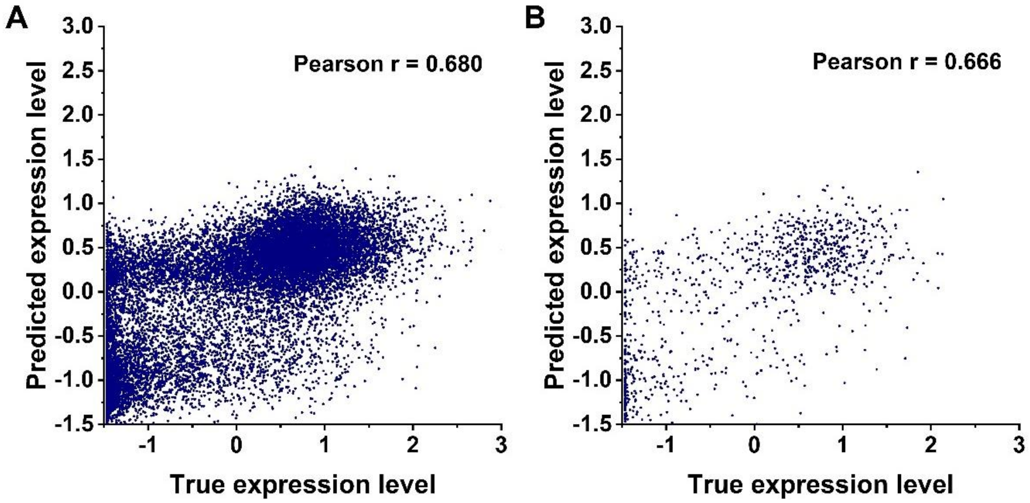


**Supplementary Fig. S3.** Prediction performance of CR-4 for (A) training dataset and (B) testing dataset.

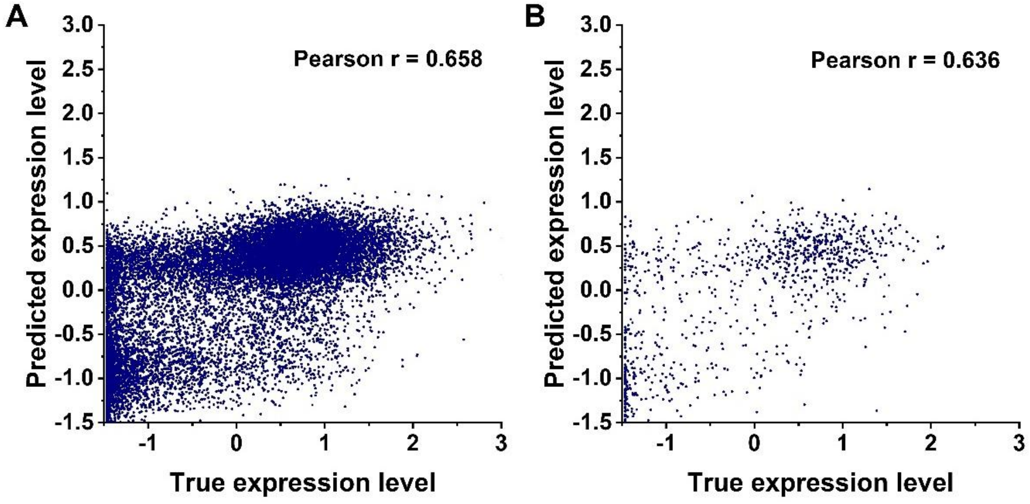


**Supplementary Fig. S4.** Prediction performance of CR-5 for (A) training dataset and (B) testing dataset.

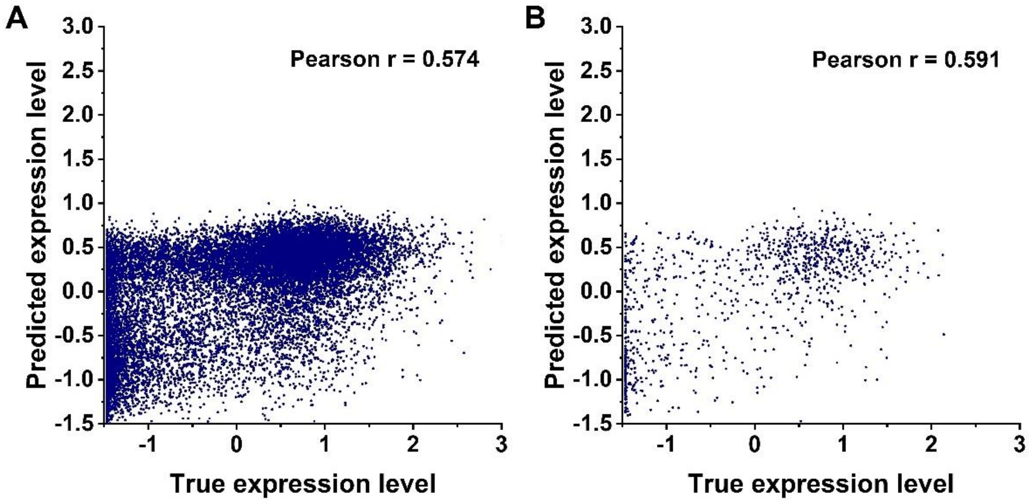


**Supplementary Fig. S5.** Prediction performance of CR-6 for (A) training dataset and (B) testing dataset.

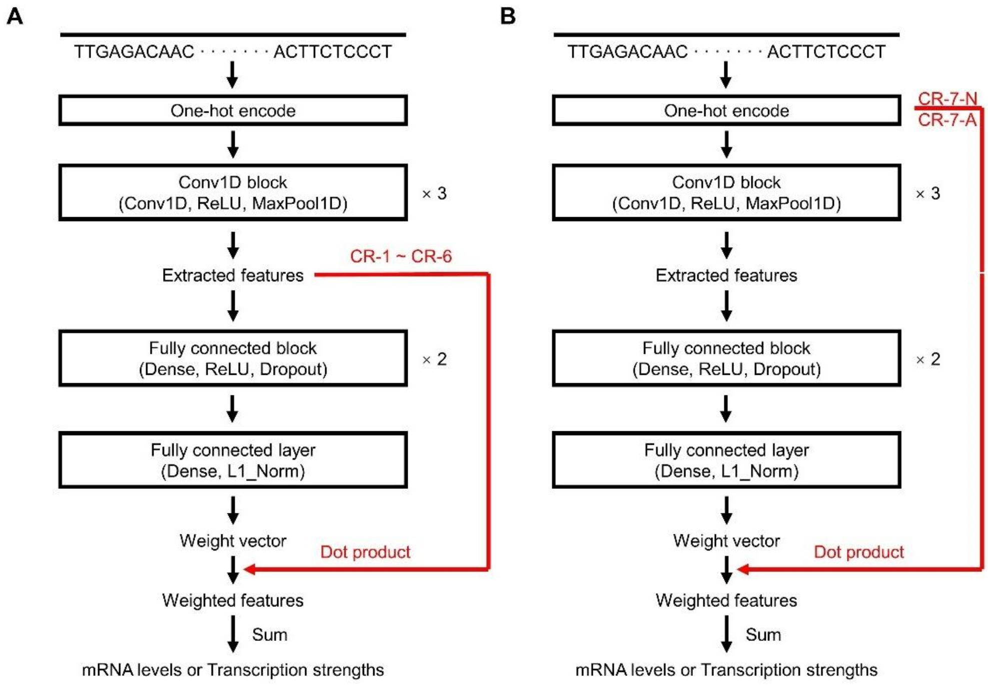


**Supplementary Fig. S6.** The comparation of the contextual regression model structures between CR-1~CR-6 and CR-7-N/A.

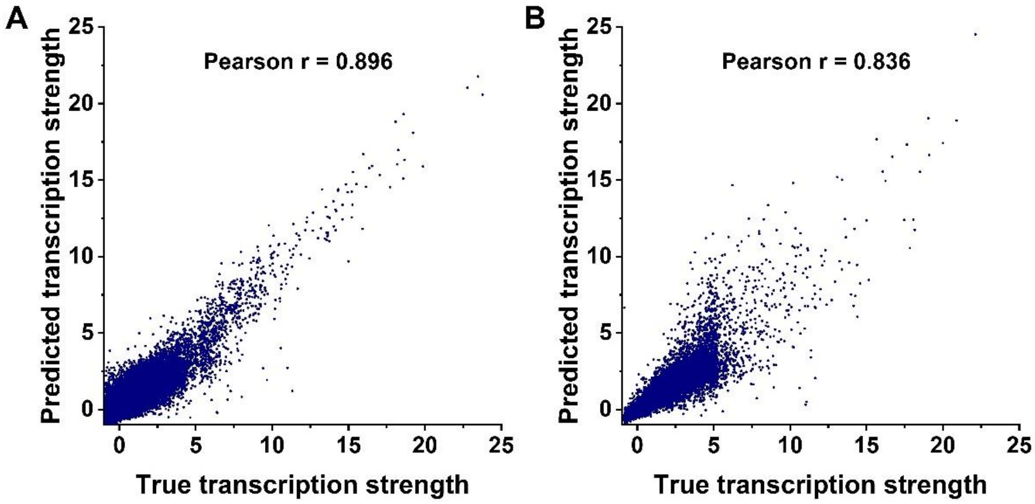


**Supplementary Fig. S7.** Prediction performance of CR-7-N for (A) training dataset and (B) testing dataset.

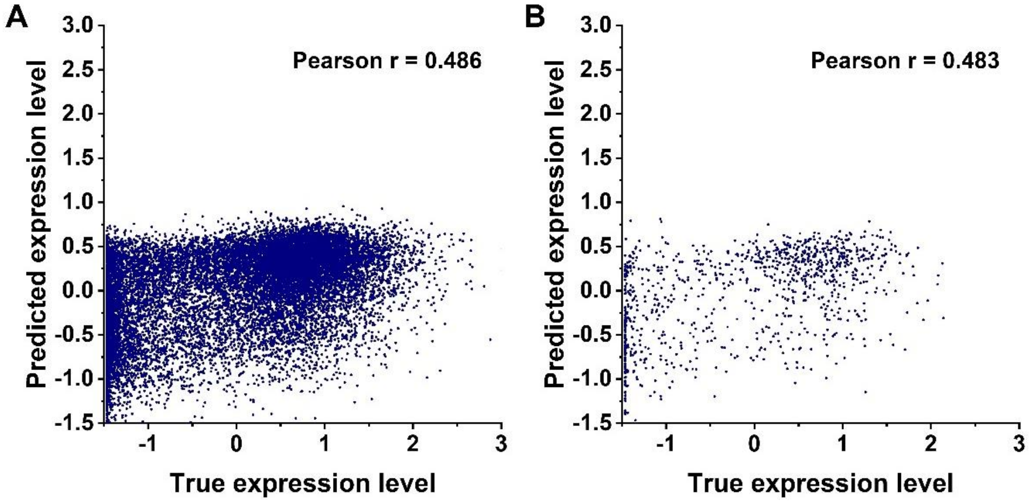


**Supplementary Fig. S8.** Prediction performance of CR-7-A for (A) training dataset and (B) testing dataset.

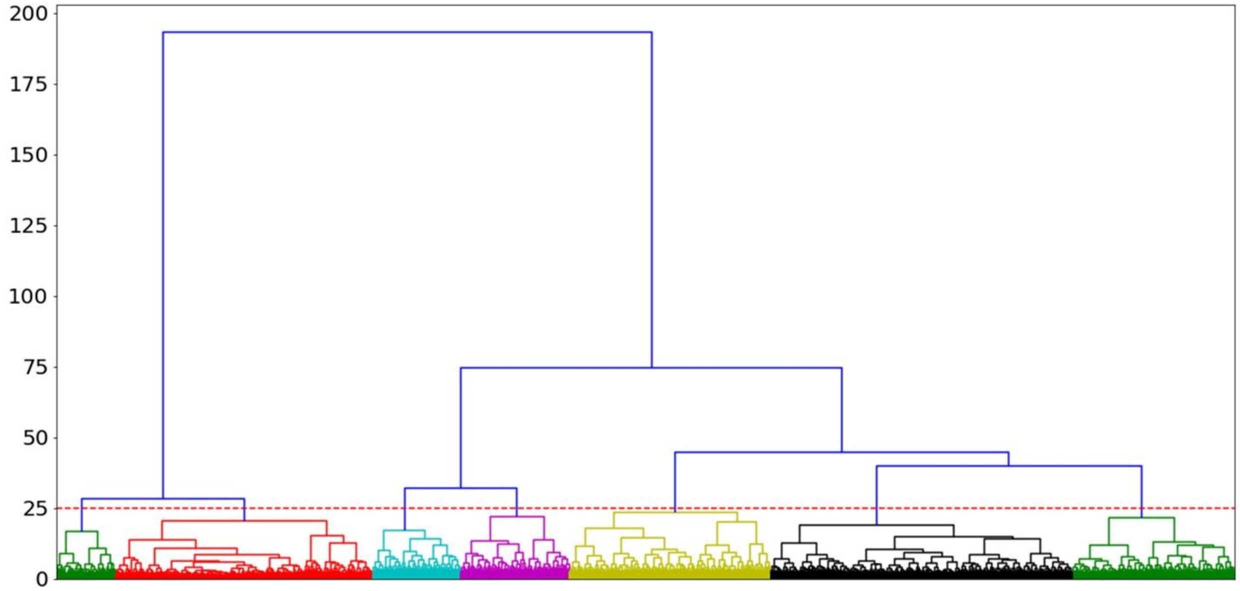


**Supplementary Fig. S9.** The hierarchical clustering of 302-bin weighted feature vectors by using the Ward variance minimization algorithm. The red dashed line is the distance threshold of 25.

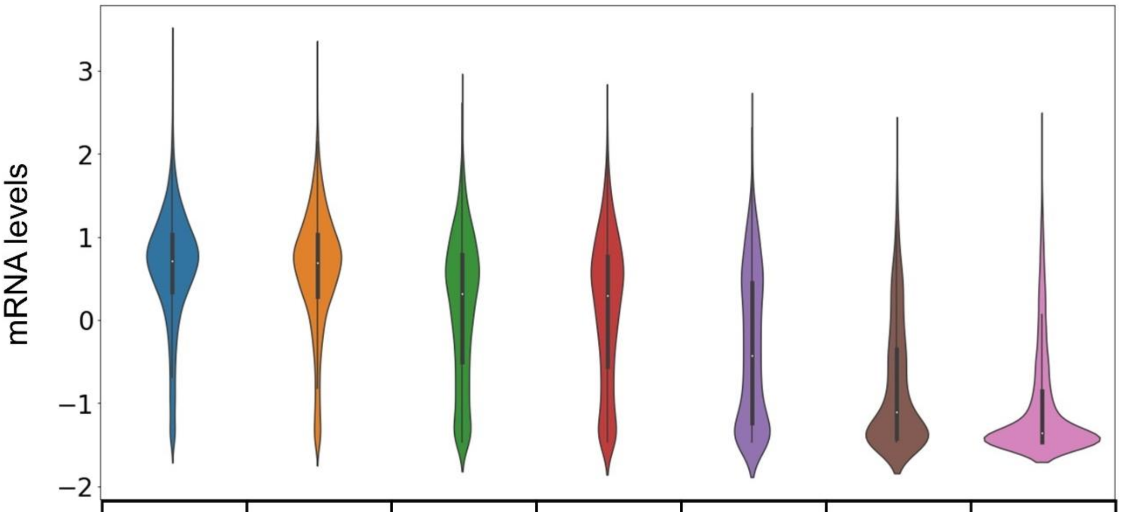


| | | | | | | | |
|---|---|---|---|---|---|---|---|
| Sample numbers | 4703 | 3136 | 2525 | 1696 | 1376 | 934 | 4007 |
| Average mRNA levels | 0.62 | 0.594 | 0.147 | 0.121 | -0.342 | -0.794 | -1.034 |

**Supplementary Fig. S10.** The distributions of mRNA levels in each group.

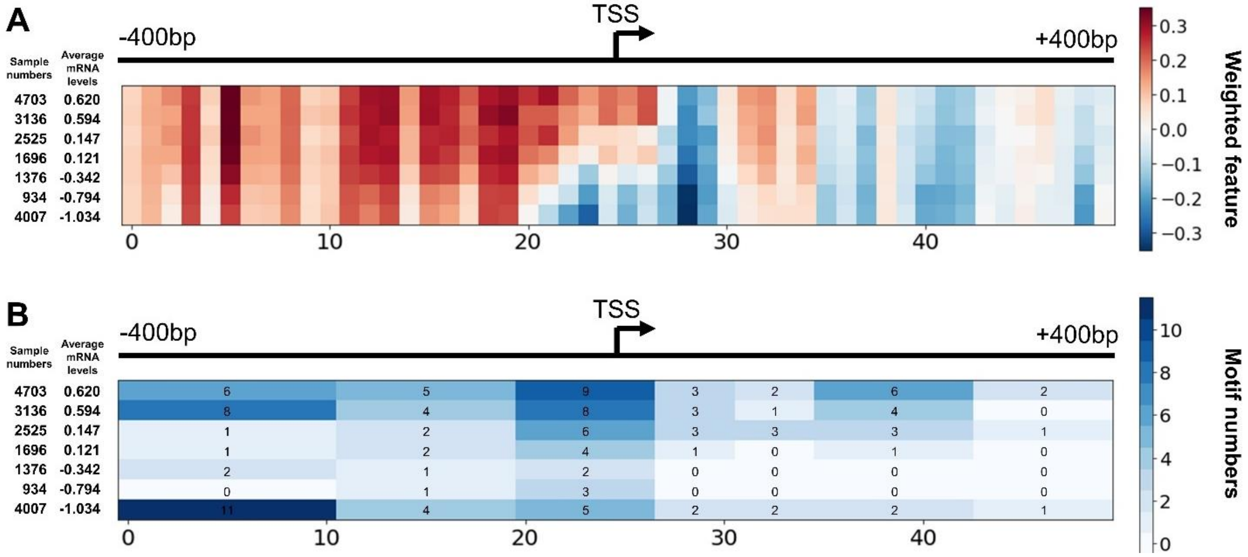


**Supplementary Fig. S11.** (A) The average weighted features of 7 groups of samples for CR-2. (B) The motif numbers in each group and each block. Left shows the average mRNA levels. The groups are ordered by the descending order of average mRNA levels.

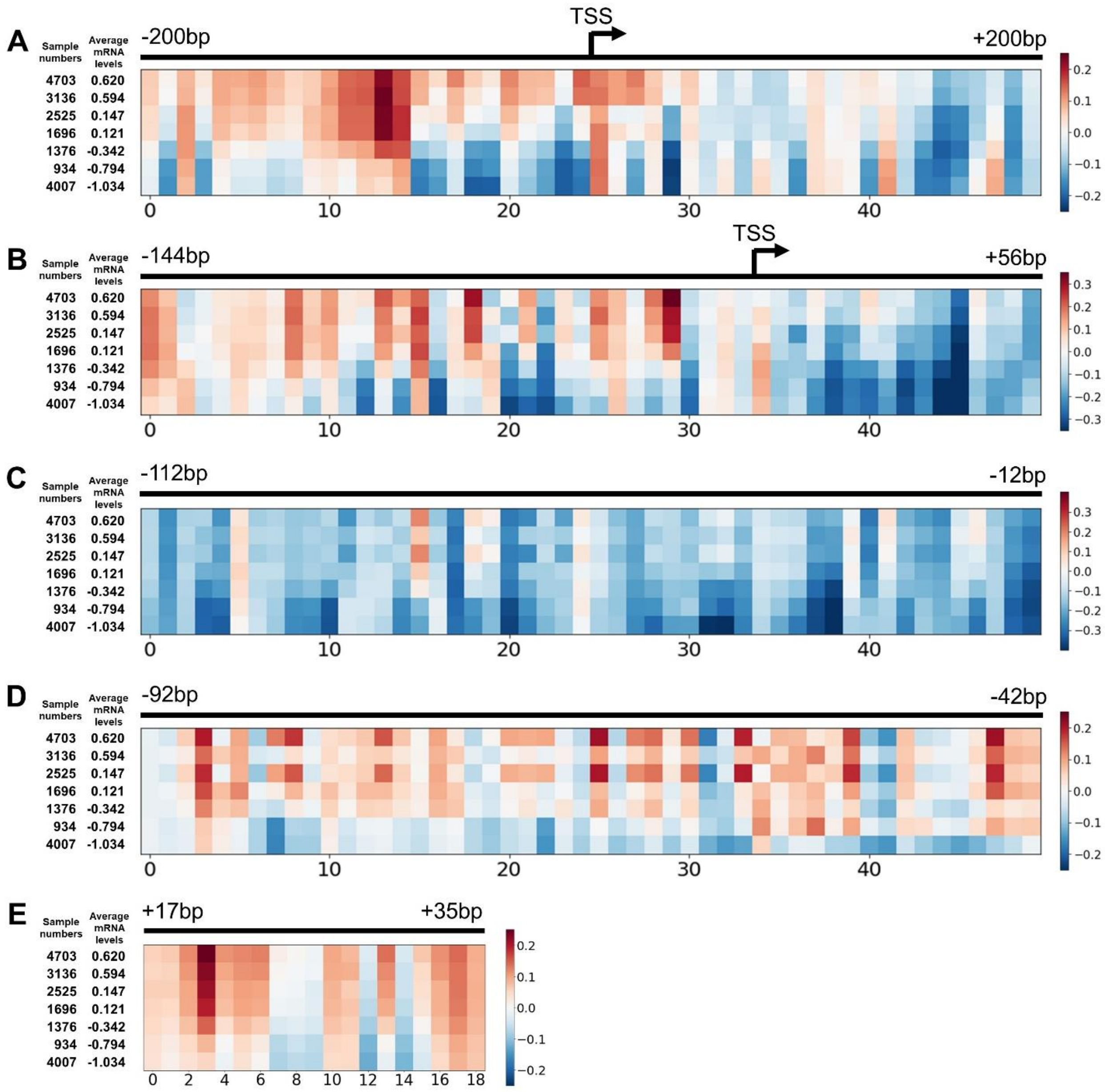


**Supplementary Fig. S12.** The average weighted features of 7 groups of samples for (A) CR-3, (B) CR-4, (C) CR-5, (D) CR-6, (E) CR-7-N.

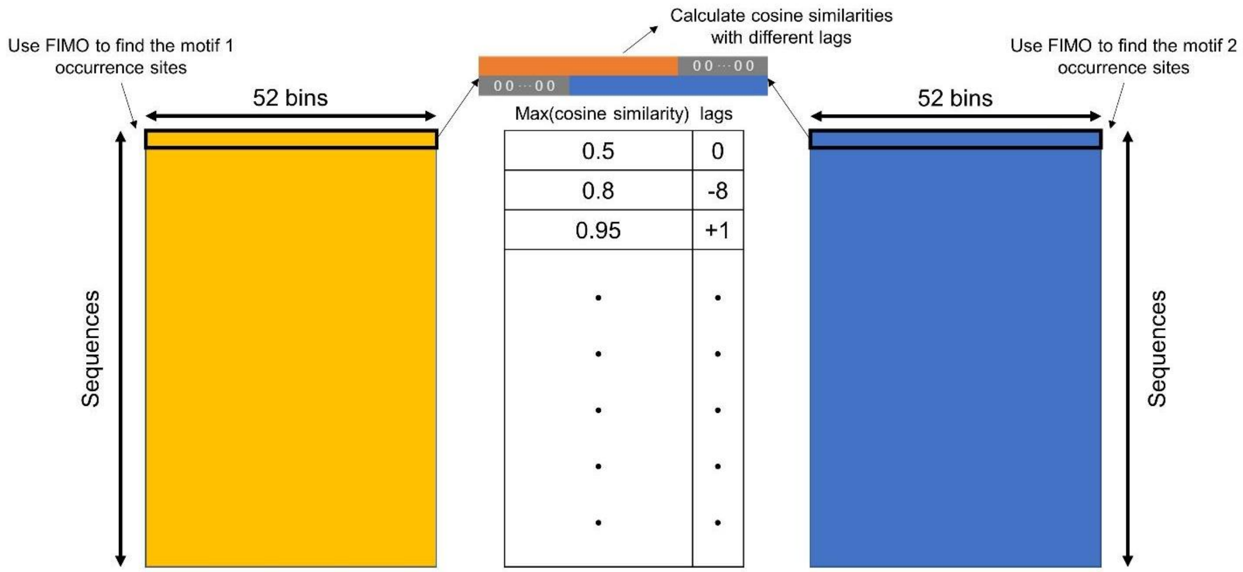


**Supplementary Fig. S13**. The process of calculating cosine similarities of occurrence sites with different lags between each pair of motifs.

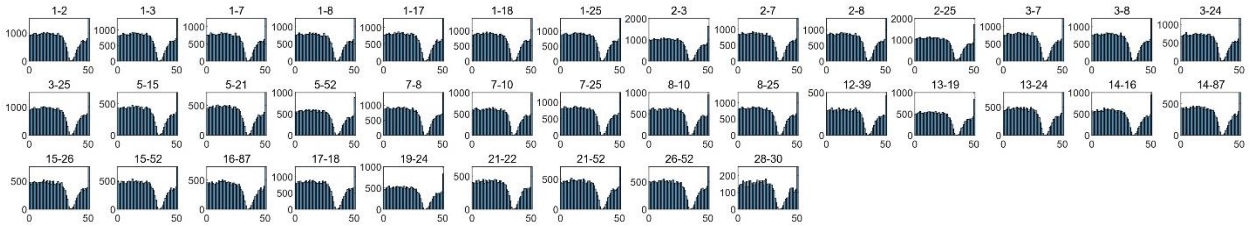


**Supplementary Fig. S14**. The distributions of the co-occurrence for the 45 motif pairs. X-axis is the bin number and the size of each bin is 200bp.

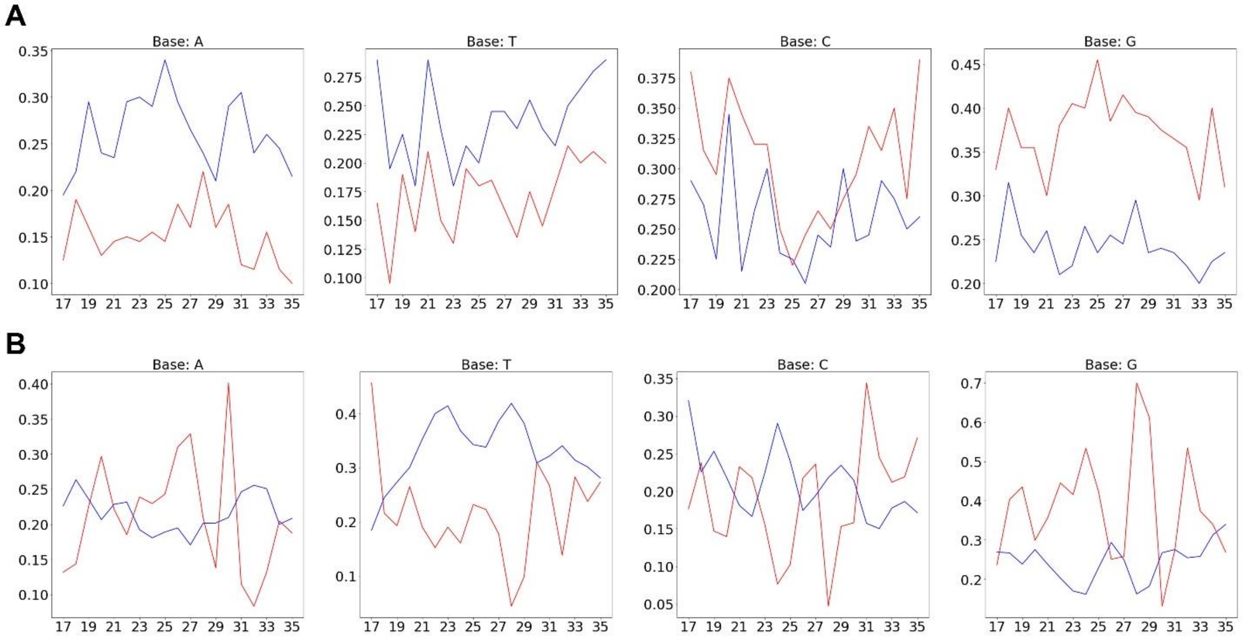


**Supplementary Fig. S15.** (A) The base frequencies in the two groups with the highest and the lowest expression levels for the genomic sequences (CR-7-N). (B) The base frequencies in two groups with the highest and the lowest expression levels for the synthetic sequences (CR-7-A). Red line represents genes with top 1% of expression levels and blue line genes with bottom 1% of gene expression levels.

**Table S1.** The results of 10 times cross-validation.

| | | 1 | 2 | 3 | 4 | 5 | 6 | 7 | 8 | 9 | 10 | Average |
|---|---|---|---|---|---|---|---|---|---|---|---|---|
| **CR-1** | Training | 0.750 | 0.761 | 0.737 | 0.755 | 0.762 | 0.748 | 0.757 | 0.749 | 0.737 | 0.748 | 0.750 |
| | Validation | 0.700 | 0.682 | 0.685 | 0.732 | 0.728 | 0.703 | 0.746 | 0.732 | 0.694 | 0.712 | 0.711 |
| **CR-2** | | 1 | 2 | 3 | 4 | 5 | 6 | 7 | 8 | 9 | 10 | Average |
| | Training | 0.709 | 0.711 | 0.715 | 0.706 | 0.719 | 0.719 | 0.707 | 0.688 | 0.714 | 0.702 | 0.709 |
| | Validation | 0.680 | 0.637 | 0.670 | 0.712 | 0.687 | 0.681 | 0.671 | 0.691 | 0.686 | 0.664 | 0.678 |
| **CR-3** | | 1 | 2 | 3 | 4 | 5 | 6 | 7 | 8 | 9 | 10 | Average |
| | Training | 0.701 | 0.698 | 0.696 | 0.691 | 0.698 | 0.699 | 0.695 | 0.694 | 0.697 | 0.717 | 0.699 |
| | Validation | 0.678 | 0.684 | 0.677 | 0.677 | 0.677 | 0.683 | 0.683 | 0.677 | 0.683 | 0.676 | 0.679 |
| **CR-1** | | 1 | 2 | 3 | 4 | 5 | 6 | 7 | 8 | 9 | 10 | Average |
| | Training | 0.674 | 0.683 | 0.688 | 0.668 | 0.682 | 0.674 | 0.667 | 0.685 | 0.681 | 0.692 | 0.679 |
| | Validation | 0.676 | 0.677 | 0.671 | 0.677 | 0.672 | 0.668 | 0.670 | 0.673 | 0.665 | 0.673 | 0.672 |
| **CR-2** | | 1 | 2 | 3 | 4 | 5 | 6 | 7 | 8 | 9 | 10 | Average |
| | Training | 0.637 | 0.645 | 0.633 | 0.635 | 0.624 | 0.653 | 0.650 | 0.629 | 0.646 | 0.631 | 0.638 |
| | Validation | 0.636 | 0.634 | 0.636 | 0.630 | 0.632 | 0.632 | 0.633 | 0.625 | 0.630 | 0.632 | 0.632 |
| **CR-3** | | 1 | 2 | 3 | 4 | 5 | 6 | 7 | 8 | 9 | 10 | Average |
| | Training | 0.590 | 0.557 | 0.586 | 0.585 | 0.591 | 0.593 | 0.578 | 0.585 | 0.573 | 0.581 | 0.582 |
| | Validation | 0.577 | 0.586 | 0.589 | 0.589 | 0.587 | 0.586 | 0.584 | 0.578 | 0.593 | 0.582 | 0.585 |
| **CR-7** | | 1 | 2 | 3 | 4 | 5 | 6 | 7 | 8 | 9 | 10 | Average |
| | Training | 0.487 | 0.497 | 0.489 | 0.508 | 0.496 | 0.496 | 0.506 | 0.483 | 0.499 | 0.499 | 0.496 |
| | Validation | 0.491 | 0.499 | 0.477 | 0.472 | 0.483 | 0.480 | 0.489 | 0.484 | 0.483 | 0.475 | 0.483 |
| **CR-8** | | 1 | 2 | 3 | 4 | 5 | 6 | 7 | 8 | 9 | 10 | Average |
| | Training | 0.900 | 0.899 | 0.890 | 0.899 | 0.903 | 0.898 | 0.888 | 0.884 | 0.900 | 0.890 | 0.895 |
| | Validation | 0.867 | 0.869 | 0.863 | 0.870 | 0.862 | 0.868 | 0.864 | 0.854 | 0.871 | 0.865 | 0.865 |

**Table S2.** Adjusting ranges of hyperparameters.

| Hyperparameter | Range |
|---|---|
| Filter size | 4~10 |
| Filter numbers | $2^2$~$2^5$ |
| Node numbers | $2^1$~$2^6$ |

**Table S3.** Comparison of prediction performance between CR-1 with Xpresso. The Xpresso results are estimated from Fig. S4 in (Agarwal and Shendure, 2020).

| | Correlation of test dataset |
|---|---|
| Xpresso, with half-life features | 0.768 |
| Xpresso, without half-life features | 0.707 |
| CR-1, without contextual regression, without half-life features | 0.727 |
| CR-1, with contextual regression, without half-life features | 0.720 |

**Table S4.** Motif Comparations with the known motifs in HOCOMOCO v11 (Kulakovskiy, *et al.*, 2018) as well as the DNA motifs that are associated with histone modifications (Ngo, *et al.*, 2019), and DNA methylation (Wang, *et al.*, 2019).

| index | query_ID | HOCOMOCO_v11 | histone | methylation |
|---|---|---|---|---|
| 1 | 'CCAGCCTGGSCRACA' | | 'H3K36me3_742' | 'MM_309.9_3.21_0.74_1_RCCTGGSTRAC' |
| 2 | 'CCTCRGCCTCCCRAR' | 'ZN770_HUMAN.H11MO.1.C' | 'H3K36me3_2886' | 'MM_814.4_2.02_0.62_8_known-PAX5' |
| 3 | 'CCTGTAATCCCAGCW' | 'PITX2_HUMAN.H11MO.0.D' | 'H3K36me3_5522' | 'MM_34.1_2.53_0.59_1_AGKCCCAGC' |
| 4 | 'CCCAGC' | | | |
| 5 | 'CAGTGAGCCGAGAT' | | 'H3K36me3_1404' | 'MM_139.5_2.02_0.61_2_GCAGTGAGC' |
| 6 | 'RGTGCRGTGGCD' | | 'H3K36me3_1507' | 'MM_412.0_2.24_0.62_3_CCATYGCAC' |
| 7 | 'GGAGRATCRCTTGAR' | | 'H3K36me3_590' | 'MM_299.9_2.20_0.63_2_ATCRCTTGAR' |
| 8 | 'AGGAGRATCRCTTGA' | | 'H3K36me3_590' | 'MM_299.9_2.20_0.63_2_ATCRCTTGAR' |
| 9 | 'CGTGATCCRCC' | | | 'MM_38.1_2.62_0.59_1_GGYGGAKCA' |
| 10 | 'GGAGAATKGCTTGA' | | | 'MM_11.7_2.05_0.54_1_AGAATGGMGHG' |
| 11 | 'TCAAAAAAAAAAAAA' | 'PRDM6_HUMAN.H11MO.0.C' | 'H3K4me1_307' | |
| 12 | 'CTAAAAATAC' | 'MEF2D_HUMAN.H11MO.0.A' | 'H3K36me3_298' | 'MM_97.6_2.15_0.57_2_GTATTTTTAG' |
| 13 | 'RGCTAATTTTTGTA' | | 'H3K36me3_186' | |
| 14 | 'AACTCCTGACCTC' | 'RARB_HUMAN.H11MO.0.D' | 'H3K36me3_3455' | |
| 15 | 'AACCCGGGAGGCGGA' | | 'H3K4me3+H3K36me3_3550' | |
| 16 | 'CAGGAGTTCRAGA' | | 'H3K36me3_800' | 'MM_11.1_2.91_0.51_1_MGGAGATCGA' |
| 17 | 'CYGTCTCWAMAAA' | | 'H3K36me3_668' | 'MM_264.2_2.04_0.62_2_GTCTCRAAM' |
| 18 | 'ARACCCYGTCTCWA' | | 'H3K36me3_1801' | 'MM_450.0_2.33_0.63_3_ACCCCGTCT' |
| 19 | 'AMAAAAATTAGCCR' | | 'H3K36me3_65' | |
| 20 | 'CAGCTAA' | | | |
| 21 | 'AGTGCARTGGYGYGA' | | 'H3K36me3_1507' | 'MM_412.0_2.24_0.62_3_CCATYGCAC' |
| 22 | 'AGTGCARTG' | | 'H3K36me3_1507' | 'MM_57.7_2.36_0.61_1_known-ZBTB3' |
| 23 | 'GCGTGAGCCACCR' | | | 'MM_306.8_2.18_0.57_4_WGGCWCACAC' |

| | | | | |
|---|---|---|---|---|
| 24 | 'CACCAYGCCYRGCTA' | | 'H3K36me3_1057' | 'MM_271.6_2.03_0.58_8_ACCVKGCCC' |
| 25 | 'AGGCTGAGGCRGGAG' | 'ZN770_HUMAN.H11MO.0.C' | 'H3K36me3_4072' | 'MM_814.4_2.02_0.62_8_known-PAX5' |
| 26 | 'AACCTCCRCCTCC' | | | 'MM_152.3_2.81_0.64_1_AASCTCCRC' |
| 27 | 'CCCGGGA' | 'ZF64A_HUMAN.H11MO.0.D' | 'H3K4me3+H3K36me3_3550' | |
| 28 | 'CTCGTGATCCRC' | | 'H3K36me3_960' | 'MM_172.5_2.58_0.57_3_ATCACGAGG' |
| 29 | 'GATCYGCCC' | | 'H3K36me3_581' | 'MM_47.1_2.01_0.56_3_GAKCTGCCY' |
| 30 | 'GACCTCGTGATCC' | 'RARA_HUMAN.H11MO.0.A' | 'H3K36me3_960' | 'MM_172.5_2.58_0.57_3_ATCACGAGG' |
| 31 | 'AGTCTCRCTC' | | | |
| 32 | 'TGGTGAAACC' | | 'H3K9me3_4143' | |
| 33 | 'ATATTTA' | | | 'MM_97.6_2.15_0.57_2_GTATTTTTAG' |
| 34 | 'AAATA' | | | 'MM_97.6_2.15_0.57_2_GTATTTTTAG' |
| 35 | 'AAATAAAT' | | | 'UM_13.5_2.17_0.53_2_DAAGTAAMTG' |
| 36 | 'AAMGAAA' | | 'H3K9me3_244' | |
| 37 | 'AAGAAAA' | | | |
| 38 | 'TATTTATTGA' | | | 'UM_10.1_2.14_0.56_1_RRTAAACAS' |
| 39 | 'TACTAAAAATA' | | 'H3K36me3_298' | 'MM_97.6_2.15_0.57_2_GTATTTTTAG' |
| 40 | 'CCCCWCCCCCNCCCC' | 'EGR1_HUMAN.H11MO.0.A' | 'H3K4me1_4240' | |
| 41 | 'GCCCCGCCCCC' | 'SP4_HUMAN.H11MO.1.A' | 'H3K4me3+H3K27ac_2294' | |
| 42 | 'CCCCGCCCCGSC' | 'KLF12_HUMAN.H11MO.0.C' | | 'UM_254.9_3.05_0.66_1_known-SP1' |
| 43 | 'CCCCRRCCCCG' | 'SP1_HUMAN.H11MO.0.A' | | |
| 44 | 'CCCCCACC' | 'TBX15_HUMAN.H11MO.0.D' | | |
| 45 | 'ACTTTGGGA' | | 'H3K36me3_4074' | 'MM_782.1_2.32_0.62_8_GCACTTTGGG' |
| 46 | 'CTCAGTTTCCTCATC' | | 'H3K4me1_4599' | |
| 47 | 'GCGGCGGCGGCGGC' | | 'H3K4me3_3087' | 'UM_3582.2_3.88_0.56_57_known-TEAD2' |
| 48 | 'CGCCGCCATCK' | 'THAP1_HUMAN.H11MO.0.C' | 'H3K4me3+H3K27ac_868' | |
| 49 | 'CGCCGCCGCCAY' | 'TAF1_HUMAN.H11MO.0.A' | 'H3K4me3_3087' | |
| 50 | 'CCTCCTCCTCCNC' | 'ZN263_HUMAN.H11MO.1.A' | 'H3K27me3_4279' | |

| | | | | |
|---|---|---|---|---|
| 51 | 'AGCCTGGG' | | 'H3K36me3_3439' | |
| 52 | 'GCTCACTGCAASCTC' | | 'H3K36me3_3613' | 'MM_139.5_2.02_0.61_2_GCAGTGAGC' |
| 53 | 'CCATCTC' | | | |
| 54 | 'AGATG' | | | |
| 55 | 'ACTACAWYTCCCAGV' | 'ZN143_HUMAN.H11MO.0.A' | 'H3K4me3_834' | |
| 56 | 'CGAGAC' | | | |
| 57 | 'CAGGAGA' | | | 'MM_323.9_2.43_0.66_2_AWYCTCCTG' |
| 58 | 'AAACCC' | | | 'MM_32.0_2.20_0.57_1_GGGGTTTCA' |
| 59 | 'ACTCACCTK' | | | |
| 60 | 'RGCCAATCRGMG' | 'NFYB_HUMAN.H11MO.0.A' | | |
| 61 | 'ACACACACACACACR' | | | |
| 62 | 'AAATGAC' | | | 'UM_23.6_2.17_0.56_2_GAAAATKAC' |
| 63 | 'CYWTTTATASM' | 'TBP_HUMAN.H11MO.0.A' | | 'UM_61.2_2.06_0.58_3_CGTTTTAHAB' |
| 64 | 'TASTA' | | | |
| 65 | 'TTTWAAA' | | 'H3K4me1+H3K36me3_306' | 'UM_61.2_2.06_0.58_3_CGTTTTAHAB' |
| 66 | 'ACGTSAC' | 'TFE3_HUMAN.H11MO.0.B' | | |
| 67 | 'KRTGAYRTCAYM' | 'ATF2_HUMAN.H11MO.0.B' | | |
| 68 | 'GAAGWCA' | | | |
| 69 | 'AGGGAAGA' | | | |
| 70 | 'CACTTCCGGBK' | 'GABPA_HUMAN.H11MO.0.A' | 'H3K4me3_902' | |
| 71 | 'CCTTCCT' | | 'H3K27me3+H3K4me1_4205' | |
| 72 | 'ACGTNG' | | | |
| 73 | 'CCCGAGTAG' | | | |
| 74 | 'AGCAGCAGCAGCAG' | 'OSR2_HUMAN.H11MO.0.C' | 'H3K4me1_2788' | 'UM_40.2_2.70_0.60_1_CGCCGCTGCTGC' |
| 75 | 'RTGGTGGYRGGYGCC' | | 'H3K36me3_3812' | |
| 76 | 'CAASATGGCBGC' | 'TYY1_HUMAN.H11MO.0.A' | 'H3K4me3+H3K27ac_868' | |
| 77 | 'CCAACATGGT' | | 'H3K36me3_1035' | 'MM_64.4_2.07_0.62_1_CCAGCATGGT' |

| | | | | |
|---|---|---|---|---|
| 78 | 'ACCATGG' | 'RFX4_HUMAN.H11MO.0.D' | | |
| 79 | 'GCGCATGCGC' | 'NRF1_HUMAN.H11MO.0.A' | 'H3K4me3+H3K27ac_1581' | |
| 80 | 'CTCCYTCCCTCCC' | 'VEZF1_HUMAN.H11MO.1.C' | | |
| 81 | 'CCCCGCGCCCCG' | 'SP2_HUMAN.H11MO.0.A' | | |
| 82 | 'CCGAGTC' | | | |
| 83 | 'CGGGGACGCGGG' | | | 'UM_91.0_3.16_0.51_3_CCGCGTCCC' |
| 84 | 'CGCTCTGTC' | | 'H3K36me3_1797' | |
| 85 | 'CTGCAGCCG' | | 'H3K4me1_3884' | |
| 86 | 'GCGCCCGGCC' | | 'H3K36me3_3401' | |
| 87 | 'AGGCTGGTCTCR' | | 'H3K36me3_1185' | |
